# BOLD signal variability and complexity in children and adolescents with and without autism spectrum disorder

**DOI:** 10.1101/491720

**Authors:** Amanda K. Easson, Anthony R. McIntosh

## Abstract

Variability of neural signaling is an important index of healthy brain functioning, as is signal complexity, which relates to information processing capacity. It is thought that alterations in variability and complexity may underlie certain brain dysfunctions. Here, resting-state fMRI was used to examine brain signal variability and complexity in male children and adolescents with and without autism spectrum disorder (ASD), a highly heterogeneous neurodevelopmental disorder. Variability was measured using the mean square successive difference (MSSD) of the time series, and complexity of these time series was assessed using sample entropy. A categorical approach was implemented to determine if the brain measures differed between diagnostic groups (ASD and typically developing (TD) groups). A dimensional approach was used to examine the continuum of relationships between each brain measure and behavioural severity, age, IQ, and the global efficiency (GE) of each participant’s structural connectome, a metric that reflects the structural capacity for information processing. Using the categorical approach, no significant group differences were found for neither MSSD nor entropy. However, the dimensional approach revealed significant positive correlations between each brain measure, GE, and age. Further, negative correlations were observed between each brain measure and behavioural severity across all participants, whereby lower MSSD and entropy were associated with more severe ASD behaviours. These results reveal the nature of variability and complexity of fMRI signals in children and adolescents with and without ASD, and highlight the importance of taking a dimensional approach when analyzing brain function in ASD.

**Significance statement:** In this study, resting-state functional magnetic resonance imaging was used to examine the variability and complexity of brain signals in children and adolescents with and without autism spectrum disorder (ASD). Group differences were not observed in variability or complexity. However, there were significant associations between these measures, behavioural severity, age, and a measure of the brain’s capacity to perform information processing, called global efficiency. Variability and complexity were higher in older participants and in participants with higher global efficiency, but were lower in participants with more severe ASD behaviours. These results provide insight into two important components of brain function in children and adolescents with and without ASD.

## Introduction

Two key components of healthy brain functioning are variability of neural signaling and complexity of these signals. In the healthy brain, variability of neural signaling allows for the formation of functional networks (Fuchs et al., 2007) and the exploration of multiple stable functional states (Ghosh et al., 2008; McIntosh et al., 2008, 2010). Previous work has shown that higher variability of brain signals is associated with better behavioural performance (Garrett et al., 2011). Further, variability changes in response to task demands: BOLD variability has been shown to be lowest in the resting-state, increased during internally focused tasks, and highest during externally focused tasks (Grady & Garrett, 2018). It has been suggested that variability allows for greater environmental uncertainty during externally-directed compared to internally-directed tasks, and variable, flexible neural signaling allows the brain to adapt to such uncertainty (Grady & Garrett, 2018).

Several studies have examined age related changes in brain signal variability. Using fMRI, Garrett et al. (2010) found that during fixation periods of a task, the standard deviation of BOLD time series increased with age in 33% of voxels and decreased in 67% of voxels. The spatial pattern of age-related changes in variability was different than that of mean BOLD activity, suggesting that these metrics provide unique information about age-related changes in brain activity. Nomi et al. (2017) examined changes in resting-state BOLD signal variability across the lifespan from ages 6 to 85 years using the mean square successive difference, and found that the majority of regions exhibited linear decreases in variability across the lifespan.

Signal complexity is also important for optimal brain function. One way to measure the complexity of a signal is with sample entropy, which assigns high values to more complex signals, and low values to highly deterministic or random signals (Costa et al., 2005). It is important to measure the complexity of brain signals in addition to variability, because a variable brain signal may not necessarily be complex. A signal with higher entropy can be interpreted as having higher information processing capacity (Heisz & McIntosh, 2013; Gatlin 1972; Shannon 1948). Like variability, complexity is thought to reflect the ability of the brain to adapt to unpredictable environments (Goldberger et al., 2002). Using fMRI, it has been shown that signal complexity in various regions of the default mode network was positively correlated with multiple cognitive functions, including attention, language, and short-term memory (Yang et al., 2013). Higher multiscale entropy of brain signals, as measured by EEG, has also been associated with greater knowledge representation (Heisz et al., 2012). Previous work using EEG has shown that entropy of brain signals increases across development (McIntosh et al., 2008; Lippé et al., 2009; Misic et al., 2010).

Abnormal levels of variability and complexity in the brain may be related to sub-optimal cognition. Too much variability can result in inefficient information processing and ineffective exploration of different network configurations in the brain (Ghosh et al., 2008; McIntosh et al., 2008, 2010). Autism spectrum disorder (ASD), a neurodevelopmental disorder that is characterized by atypical social communication and restricted, repetitive and stereotyped behaviours (American Psychiatric Association, 2013), is one condition that may be characterized by detrimental levels of noise (Rubenstein & Merzenich, 2003). Studies of dynamic FC have shown that FC is also more variable in ASD (e.g. Falahpour et al., 2016; Zhang et al., 2016). However, the nature of resting-state BOLD signal variability in ASD compared to typically developing (TD) individuals remains unclear. Entropy of EEG signals has also been shown to be reduced in ASD at rest (Bosl et al., 2011) and during both social and non-social tasks (Catarino et al., 2011), but the nature of the complexity of resting-state BOLD time series in ASD is currently unknown.

In the present study, we characterized BOLD signal variability and complexity in youth with and without ASD. Further, we characterized relationships between these measures and age, structural connectivity, and behavioural severity.

## Materials and Methods

### Participants

Twenty male participants with ASD (M = 13.25 years, SD = 2.87 years) and 17 male TD participants (M = 13.42 years, SD = 3.21 years) from the San Diego State University sample from the Autism Brain Imaging Data Exchange (ABIDE) II database (Di Martino et al., 2017) were included in this study. Informed consent or assent had been obtained for all participants and caregivers in accordance with the University of California, San Diego and San Diego State University Institutional Review Boards. Participants were matched for age, full-scale IQ and head motion (Table 1). Participants were excluded if they were less than 8 years of age, their full-scale IQ was below 75 or if their head motion during the scan exceeded a mean framewise displacement 0.2mm. Further, participants were included only if they had an MPRAGE scan, resting-state fMRI scan, and DWI scan.

**Table 1:** Participant Characteristics

| **Parameter** | **ASD**<br>**(Mean ± SD)**<br>**[range]** | **TD**<br>**(Mean ± SD)**<br>**[range]** | **Group difference** |
| --- | --- | --- | --- |
| N | 20 | 17 | N/A |
| Age | 13.25 ± 2.87<br>[9.6 – 17.80] | 13.42 ± 3.21<br>[8.10 – 17.60] | $t(35) = -0.17, p = 0.87$ |
| IQ | 98.50 ± 14.33<br>[77 – 130] | 103.35 ± 10.15<br>[79 – 125] | $t(35) = -1.17, p = 0.25$ |
| Handedness | 15 RH<br>3 LH<br>2 mixed | 14 RH<br>1 LH<br>2 mixed | $X^2(2, N=37) = 0.80, p = 0.67$ |
| Head motion (mean<br>FD) | 0.106 ± 0.035<br>[0.054 – 0.189] | 0.096 ± 0.042<br>[0.040 – 0.194] | $t(35) = 0.76, p = 0.45$ |
| SRS Total | 104.25 ± 24.92<br>[59 – 147] | 18.41 ± 11.57<br>[2 – 36] | $t(35) = 13.04, p < 0.001$ |
| SRS Awareness | 13.10 ± 4.39<br>[3 – 19] | 3.82 ± 2.83<br>[0 – 10] | $t(35) = 7.48, p < 0.001$ |
| SRS Cognition | 17.85 ± 6.02<br>[4 – 28] | 2.94 ± 2.77<br>[0 – 8] | $t(35) = 9.39, p < 0.001$ |
| SRS Communication | 34.95 ± 10.36<br>[21 – 55] | 5.88 ± 4.48<br>[0 – 15] | $t(35) = 10.72, p < 0.001$ |
| SRS Motivation | 17.80 ± 4.12<br>[11 – 25] | 3.94 ± 2.44<br>[0 – 8] | $t(35) = 12.15, p < 0.001$ |
| SRS Mannerisms | 22.05 ± 7.86<br>[8 – 36] | 1.82 ± 2.77<br>[0 – 11] | $t(35) = 10.08, p < 0.001$ |
| SCQ | 20 ± 5.82<br>[3 – 30] | 2.42 ± 1.97<br>[0 – 7] | $t(35) = 11.87, p < 0.001$ |
| ADOS2 Total | $14.70 \pm 4.92$<br>[5 – 24] | N/A | N/A |
| ADOS2 SA | $11.35 \pm 4.26$<br>[5 – 20] | N/A | N/A |
| ADOS2 RRB | $3.35 \pm 1.95$<br>[0 – 8] | N/A | N/A |
| ADOS2 Severity | $7.85 \pm 2.08$<br>[3 – 10] | N/A | N/A |

### Data acquisition

The following scans were acquired on a GE MR750 system at SDSU: structural (T1-weighted 3D SPGR sequence; TR = 8.136 ms, TE = 3.172 ms, TI = 600 ms, flip angle = 8°, field of view (FOV) = 256 mm, matrix size 256 × 192, 1.0 mm isotropic voxel resolution), diffusion-weighted (T2-weighted sequence; TR = 8500 ms, FOV = 192 mm, matrix size 96 × 96, in-plane voxel dimension of 0.9375 × 0.9375mm/1.875 × 1.875mm with 2mm slice thickness, 68 slices, 61 diffusion directions), and fMRI (two-dimensional T2-weighted gradient echo planar imaging blood oxygen level-dependent contrast sequence; TR = 2000 ms, TE = 30 ms, flip angle = 90°, FOV = 200 mm, in-plane voxel dimension of 3.4375 mm x 3.4375 mm voxel resolution, 3.4 mm slice thickness, matrix size 64 × 64, 42 slices, 180 TRs, eyes open).

### Resting-state fMRI preprocessing

Data were preprocessed using the Optimization of Preprocessing Pipelines for NeuroImaging (OPPNI) software (Churchill et al., 2012a, 2012b, 2015). First, motion correction was performed using AFNI’s 3dvolreg function, followed by the generation of subject-specific non-neuronal tissue masks using the PHYCAA+ algorithm (49), replacement of outlier volumes with interpolated values from neighbouring time points (Campbell et al. (2013); https://www.nitrc.org/projects/spikecor_fmri), slice-timing correction, and temporal detrending using a second-order Legendre polynomial. Next, principal component analysis was performed on the motion parameters obtained from the motion correction step. Principal components that accounted for more than 85% of the variance of the motion parameters were regressed out of the fMRI data. The time series of the mean white matter and CSF signals were regressed out of the data, and finally, lowpass filtering was performed with a cutoff of 0.1Hz.

The ROI atlas used in this study is described in Bezgin et al. (2012). Parcellation of each hemisphere into 48 regions was performed on a monkey brain surface using mapping rules established by Kotter and Wanke (2005). The parcellation was then transformed to the MNI human brain using landmarks from Caret (www.nitrc.org/projects/caret; Van Essen et al., 2001; Van Essen & Dierker, 2007). For this study, we only used cortical regions from the atlas, resulting in 82 ROIs in total (41 per hemisphere; Table 2; Figure 1). The atlas was transformed from MNI space to each subject’s T1 space, then subsequently transformed to each subject’s functional space using the inverse of each subject’s functional-to-anatomical transform, which was obtained using linear registration with 6 degrees of freedom (DOF). Next, the time series of each region was extracted using the mri_segstats function in Freesurfer.

**Table 2:**
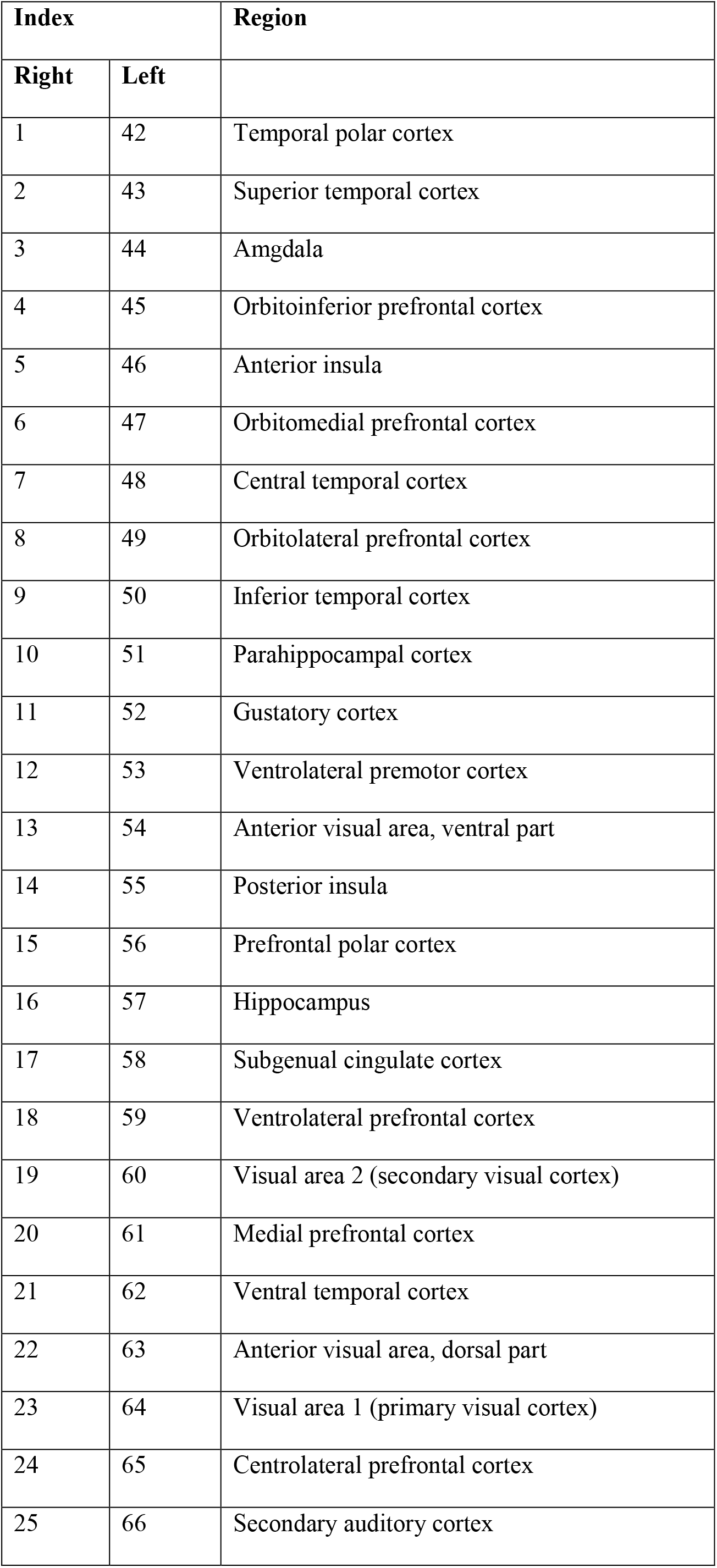
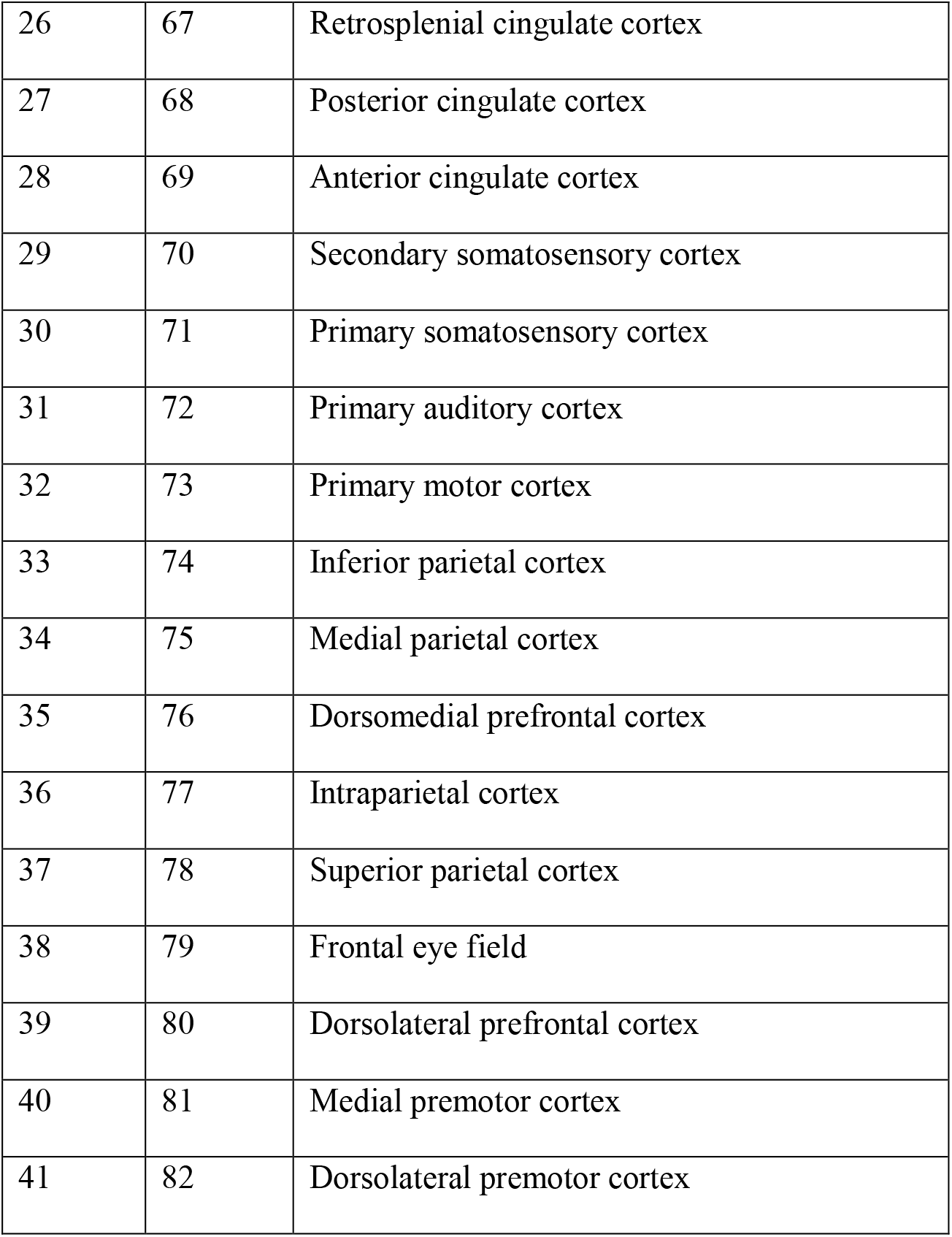
Cortical regions from the Bezgin et al. (2012) atlas

**Figure 1:**
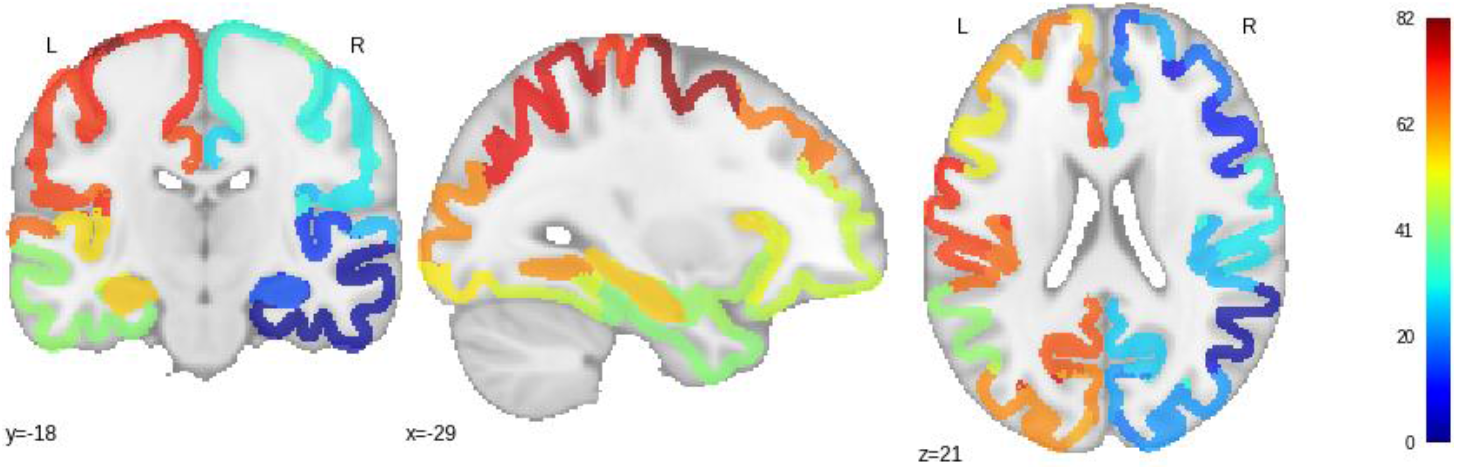
Cortical regions from the Bezgin et al. (2012) atlas in MNI space

### DWI preprocessing

Preprocessing of dwMRI data included motion correction using eddy current correction in FSL, fitting diffusion tensor models at each voxel using the dtifit function in FSL, co-registering each subject’s skull-stripped T1 MRI to DTI space using linear registration with 12 degrees of freedom (DOF), registering a standard MNI T1 image to each subject’s T1 image, labeling the grey matter in T1 space with the 82 cortical regions using nearest neighbour interpolation, then transforming the ROI-labelled T1 image into diffusion space. BEDPOSTX (Bayesian Estimation of Diffusion Parameters Obtained using Sampling Techniques; the X refers to the modeling of crossing fibres) in FSL was used to fit the probabilistic diffusion model on the preprocessed dwMRI data. Markov Chain Monte Carlo sampling is run to build up distributions on diffusion parameters at each voxel. Probabilistic fiber tracking using the probtrackx2 function in FSL was then performed to define weights (fiber counts/number of streamlines) and anatomical distances between each pair of ROIs. Probtrackx2 takes repetitive samples from the distributions of voxel-wise principal diffusion directions. A streamline is computed through each of these local samples, thus generating a probabilistic streamline. By taking many samples, a histogram of the posterior distribution of the streamline location, or connectivity distribution, is created. All masks for tractography were interface masks, that is, masks of the boundary between the gray matter and white matter. This approach is referred to as anatomically-constrained tractography, and allows one to avoid the white matter seeding bias, that is, the tendency for major white matter structures to be over-defined (Smith et al., 2012). The seeds for tractography were masks of each of the 82 ROIs. The following settings were used: 5000 samples, 2000 steps per sample, step length of 0.5, curvature threshold of 0.2, loop check on, and correction of the path distribution for the length of the pathways. Targets were masks for each of the other 81 ROIs; streamlines were terminated once they reached the target mask. For a given ROI, the exclusion mask for connections with ipsilateral ROIs consisted of that ROI and ROIs in the contralateral hemisphere, and the exclusion mask for connections with contralateral ROIs was the mask for that ROI. Any pathways that entered the exclusion masks were discarded. For each subject, an SC weights matrix was defined for each pair of ROIs as the number of streamlines detected between the two ROIs divided by the total number of samples. A tract lengths matrix was defined for each pair of ROIs as the average distance between the two ROIs. Finally, both the weights and tract lengths matrices were thresholded based on tractography data for the same atlas from the CoCoMac database (Stephan et al., 2001), such that non-existent connections in the CoCoMac data were set to 0 in the human weights and tract lengths matrices to control for false positives.

Next, the global efficiency (GE) of each participant’s SC weights matrix was calculated. GE was used as a summary measure of structural networks because a single value is calculated for the entire network, instead of one measure for each ROI. GE is defined as the average inverse shortest path length in the brain network, and is a measure of the overall capacity of the brain network to perform “parallel information transfer and integrated processing” (Bullmore & Sporns, 2012). Therefore, it would be expected that brain networks that exhibit high GE in their structural networks would exhibit high overall entropy of functional networks. GE was calculated for each participant’s structural connectivity matrix using the efficiency_wei.m function from the Brain Connectivity Toolbox (Rubinov & Sporns, 2010).

### Experimental Design and Statistical Analysis

#### BOLD signal variability and complexity

BOLD signal variability was defined using the mean square successive difference (MSSD), which is defined as:

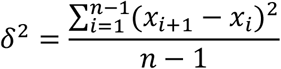

Complexity was defined as the sample entropy of the signal using the sample entropy function from PhysioNet (https://physionet.org/physiotools/mse/). Sample entropy is used to determine the appearance of repetitive patterns in a time series, thus measuring the regularity of the time series. As such, sample entropy values are low for signals that are completely random or strongly deterministic, and are high for more complex signals. (Costa et al., 2005; Pincus 1991; Richman & Moorman, 2000).

One issue with measuring sample entropy is that two parameters must be selected: the pattern length *m*, which is the number of data points used for pattern matching, and the tolerance factor *r*, which is the fraction of the standard deviation of the signal. Recently, Yang et al. (2018) presented a method for selecting optimal m and r values for an fMRI dataset based on the standard error of entropy estimates in CSF voxels. For different combinations of m and r, entropy was calculated for each voxel in each participant’s CSF mask. Then, the standard error of this entropy estimate was defined as

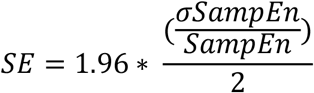

as in Yang et al. (2018). The median standard error was then calculated across participants, for each combination of m and r. Further, the number of sample entropy estimation failures due to an absence of pattern matches was summed across all CSF voxels and participants. “Acceptable” combinations of m and r were defined as those with a median standard error of less than 0.1, and with zero sample entropy estimation failures across all CSF voxels and participants. These values were used to calculate the sample entropy of the ROIs in the grey matter. Subsequently, for each participant, sample entropy values were averaged across the acceptable m and r combinations for each ROI. Using this method, we tested m values of 1 to 4 with a step size of 1, and r values ranging from 0.05 to 0.80, with a step size of 0.05, which were the same r values used in Yang et al. (2018). These investigators also tested greater values of m, but found that for m > 4, the standard error was greater than 0.1 for all values of r; thus, we only test m values of 4 or less.

### Partial least squares

Partial least squares (PLS; McIntosh et al., 1996; McIntosh & Lobaugh, 2004) is a multivariate statistical method that is used to determine optimal relationships between a set of brain variables and either a study design (mean-centering PLS) or a set of behaviour variables (behavioural PLS). In PLS, singular value decomposition is used to calculate orthogonal patterns that explain the maximal covariance between brain variables and design or behaviour variables. “Brain saliences” show which brain variables best characterize the relationship between brain and design/behaviour. In mean-centering PLS, design saliences indicate the group, condition, or group *x* condition profiles that best characterize the relationship between the brain and design variables. In the case of behavioural PLS, behaviour saliences indicate the profile of behaviour scores that best characterize the relationship between the set of brain variables and behaviour variables. Further, singular values show the proportion of covariance that each pattern (relationship between brain and design/behaviour) accounts for.

In this study, mean-centering PLS was used to determine optimal contrasts in entropy between conditions (acceptable combinations of m and r), and optimal contrasts in MSSD and entropy between ASD and TD groups. Behavioural PLS was used to determine relationships each of these brain variables and a set of predictor variables, including GE of the structural networks, age, IQ, scores on the Social Responsiveness Scale (SRS; Constantino & Gruber, 2005), and scores on the Social Communication Questionnaire (SCQ; Rutter et al., 2003). The SRS is a parent or teacher report of ASD-related traits that was created for use in the general population in both clinical and educational settings. The SCQ is a caregiver report that is used as a screening measure for ASD and was developed based on the Autism Diagnostic Interview-Revised (Lord et al., 1994). Behavioural PLS was also performed using scores on the Autism Diagnostic Observation Schedule 2 (ADOS2) instead of SRS and SCQ scores in the ASD group. This analysis was only performed for ASD participants, as ADOS2 scores were not available for the TD group.

The significance of each PLS pattern can be determined using permutation testing. The rows (participants) of the data matrix were reshuffled and the singular value was recalculated. This procedure was repeated 1000 times to obtain a distribution of singular values. Then, a p-value for the original singular value was obtained by calculating the proportion of singular values from the sampling distribution that are greater than the original singular value. Further, the reliability of each brain salience can be determined using a bootstrapping procedure. Here, 500 bootstrap samples were generated by randomly sampling participants with replacement while maintaining group membership. Next, a bootstrap ratio (BSR) was calculated as the ratio of the brain salience to the standard error of the salience from the bootstrap samples, which is a measure of the stability of the salience regardless of which participants are included in the analysis. Stable brain regions were defined as those that surpassed a BSR threshold of +2, which corresponds to approximately a 95% confidence interval.

### Data visualization

Figures were created using Matlab (MATLAB 8.6.0 (R2015b), MathWorks, Natick, MA). Violin plots were created using the distributionPlot.m function (Jonas 2017).

## Results

### Effects of m and r on entropy estimates

Figure 2 shows the results of the entropy estimates in CSF voxels. This analysis revealed that for m = 1, all r values were acceptable. For m = 2, r values from 0.20 to 0.65 were acceptable. Hereafter, the term “condition” will be used to refer to an acceptable combination of m and r.

**Figure 2:**
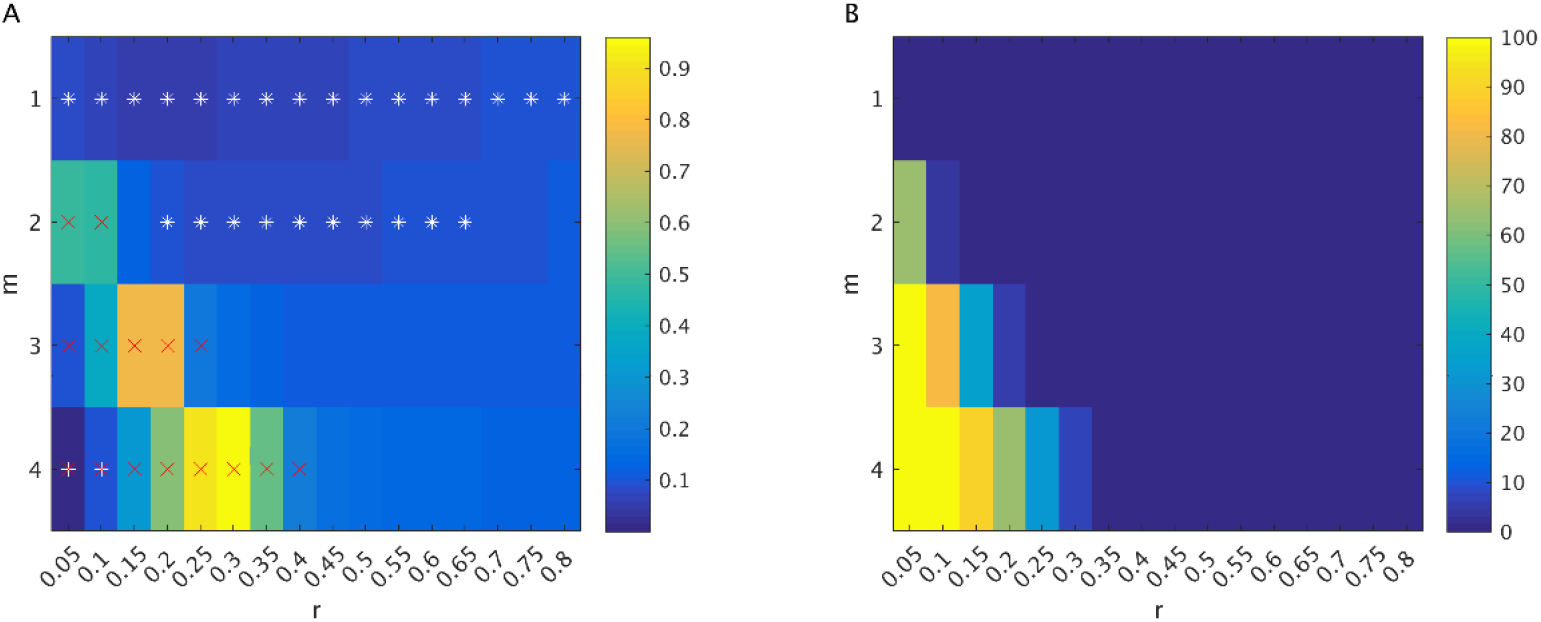
A) Standard error of entropy estimates in CSF voxels. White stars indicate “acceptable” combination of m and r, for which the median entropy estimate in CSF voxels across participants was less than 0.1. Red crosses indicate combinations of m and r that resulted in a failure of entropy estimation in at least 1 voxel across all participants. B) Percentage of all CSF voxels (across all participants) showing a failure of entropy estimation.

To ensure that any group differences in MSSD and entropy in the gray matter ROIs were not confounded by residual effects of head motion, we first performed a behavioural PLS using mean framewise displacement (FD) as the “behaviour” variable. We found that there was no significant relationship with head motion for MSSD (*p* = 0.53) or entropy (*p* = 0.23). However, we found that the relationship between SD and head motion was significant (*p* = 0.002), even when additional ICA denoising was implemented using ICA-AROMA (*p* = 0.005; Pruim et al., 2015a, 2015b). These findings therefore provide more support for the use of MSSD over SD for measuring brain signal variability.

Mean-centering PLS was then used to examine how different acceptable combinations of m and r affected entropy estimates in the 82 grey matter ROIs across all participants. There was one significant LV from this analysis that showed a contrast between different combinations of m and r (p < 0.001, 99.97% covariance explained; Figure 3). Therefore, entropy estimates differ significantly depending on which combinations of m and r are used for the analyses.

**Figure 3:**
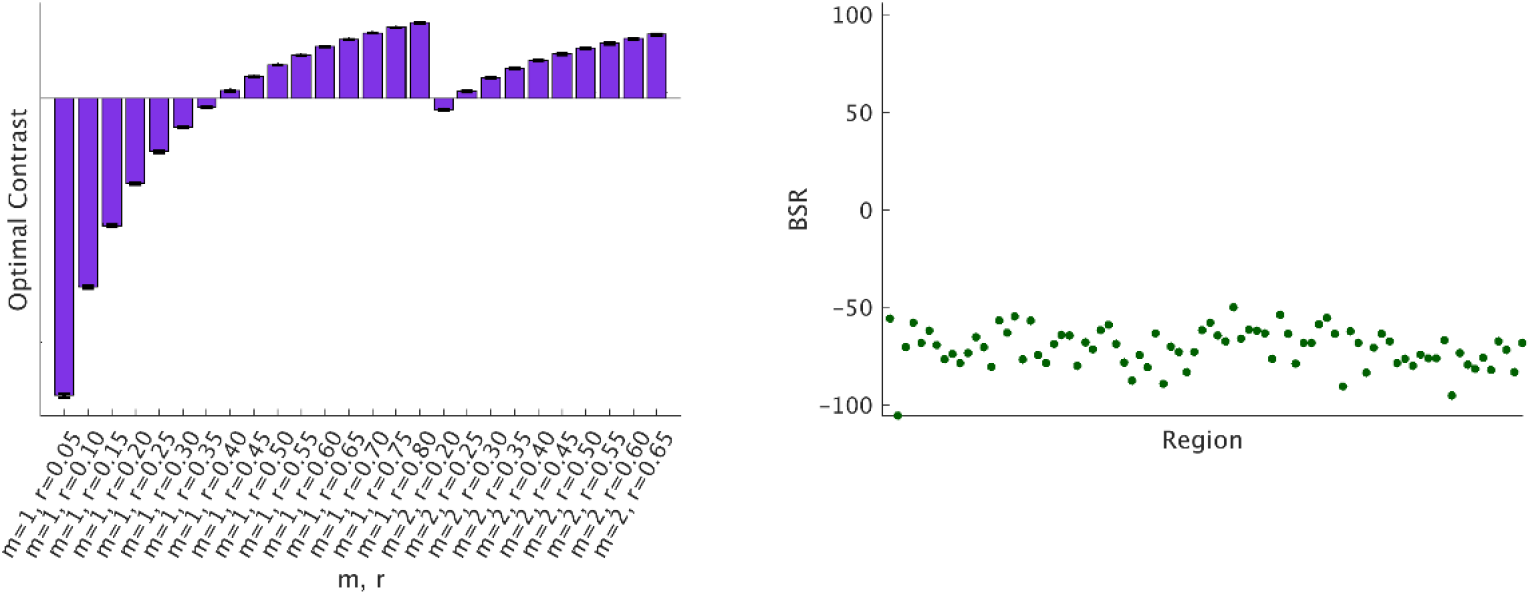
Contrast between entropy conditions and associated BSRs for each region, at a threshold of +2. Error bars show 95% confidence intervals determined through bootstrap resampling.

For subsequent analyses, since there were differences in entropy that depended on the choice of m and r, we averaged entropy estimates across all acceptable m = 2 conditions except for r = 0.20, which showed a contrast with the other r values for m = 2.

### Categorical analyses of MSSD and entropy using mean-centering PLS

Next, mean-centering PLS was performed to determine differences in MSSD and entropy between diagnostic groups. In other words, a categorical approach was used to analyze MSSD and entropy. For MSSD, no significant differences were observed between ASD and TD groups (*p* = 0.18). Similarly, groups were not significantly different in entropy (*p* = 0.15).

### Dimensional analyses of MSSD and entropy using behavioural PLS

Next, behavioural PLS was performed to characterize relationships between MSSD and entropy in each brain region and the following measures: GE, age, IQ, SRS scores, and SCQ scores and between entropy and these measures. One main advantage of using PLS in this way is that relationships between different “behaviour” variables in relation to brain variables can be analyzed, for instance, to determine if two behaviour variables differ in terms of the strength or direction of their linear relationships with the brain variables.

Prior to performing the PLS analysis, the relationships between the set of behaviour variables were analyzed. As shown in Figure 4, there was a moderate positive correlation between GE and age, a weak negative correlation between GE and IQ, and a moderate negative correlation between GE and SRS and GE and SCQ scores. SRS and SCQ scores were strongly positively correlated with each other, and weakly negatively correlated with IQ. Further, GE did not differ significantly between the ASD and TD groups, *t*(35) = −1.58, *p* = 0.12.

**Figure 4:**
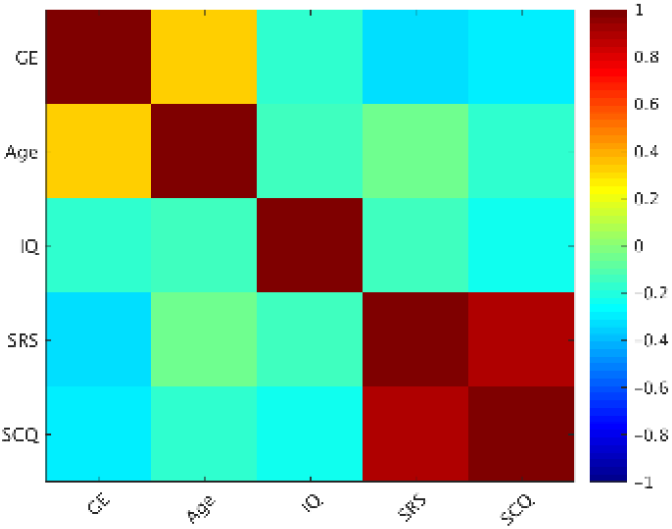
Correlation matrix showing relationships between the set of predictor variables used in the behavioural PLS for all participants.

The behavioural PLS analyses revealed one significant LV for MSSD (*p* = 0.004, 68.46% covariance explained, Figure 5) and for one significant LV for entropy (*p* = 0.008, 57.56% covariance explained, Figure 6). For both analyses, there was a distributed set of brain regions that exhibited positive correlations with GE and IQ, but negative correlations with the SRS and SCQ measures. For both MSSD and entropy, a continuum of brain-behaviour relationships can be observed from ASD to TD participants, as shown in the middle panel in Figures 5 and 6. In other words, a broad range of brain and behaviour scores can be observed across all participants, but overall, brain and behaviour scores were higher in the TD group. Further, the brain saliences for the MSSD and entropy analyses were significantly correlated, *r* = 0.47, *p* = 0.002 (1000 permutations), showing that there was a strong relationship between the brain regions that contributed to the MSSD pattern and the regions that contributed to the entropy pattern (Figure 7).

**Figure 5:**
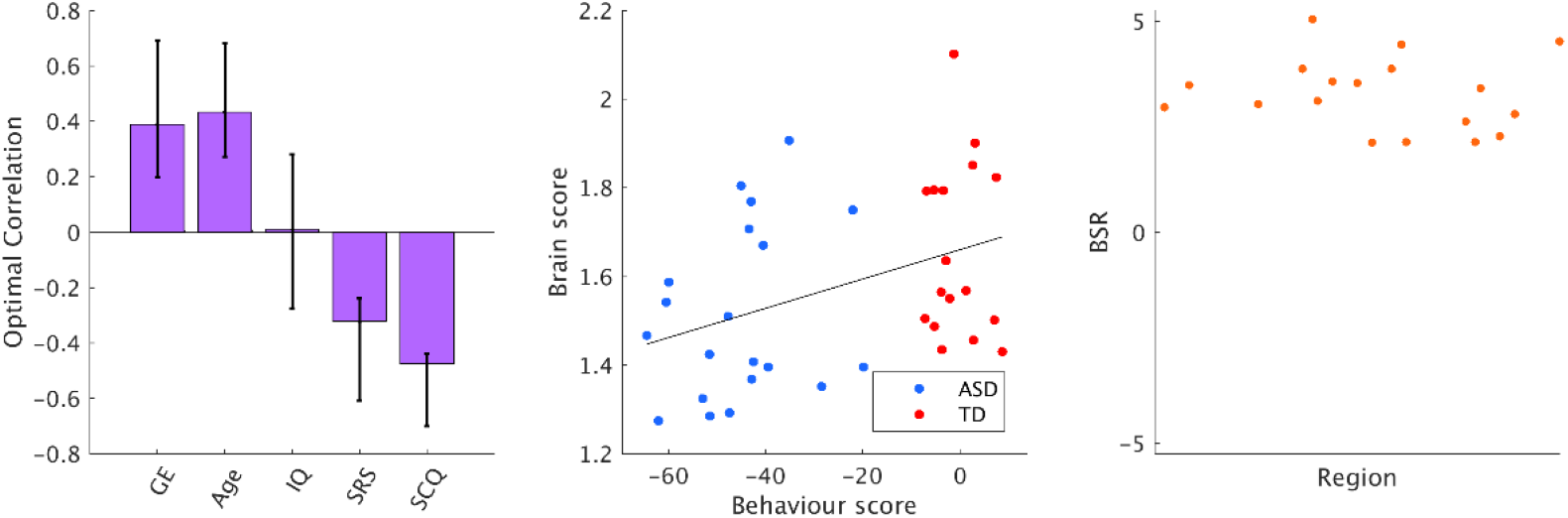
Contrast in relationships for correlations between MSSD and predictor variables, associated brain and behaviour scores for each group, and BSRs for each region, at a threshold of +2. Error bars show 95% confidence intervals determined through bootstrap resampling. Blue circles = ASD, red circles = TD.

**Figure 6:**
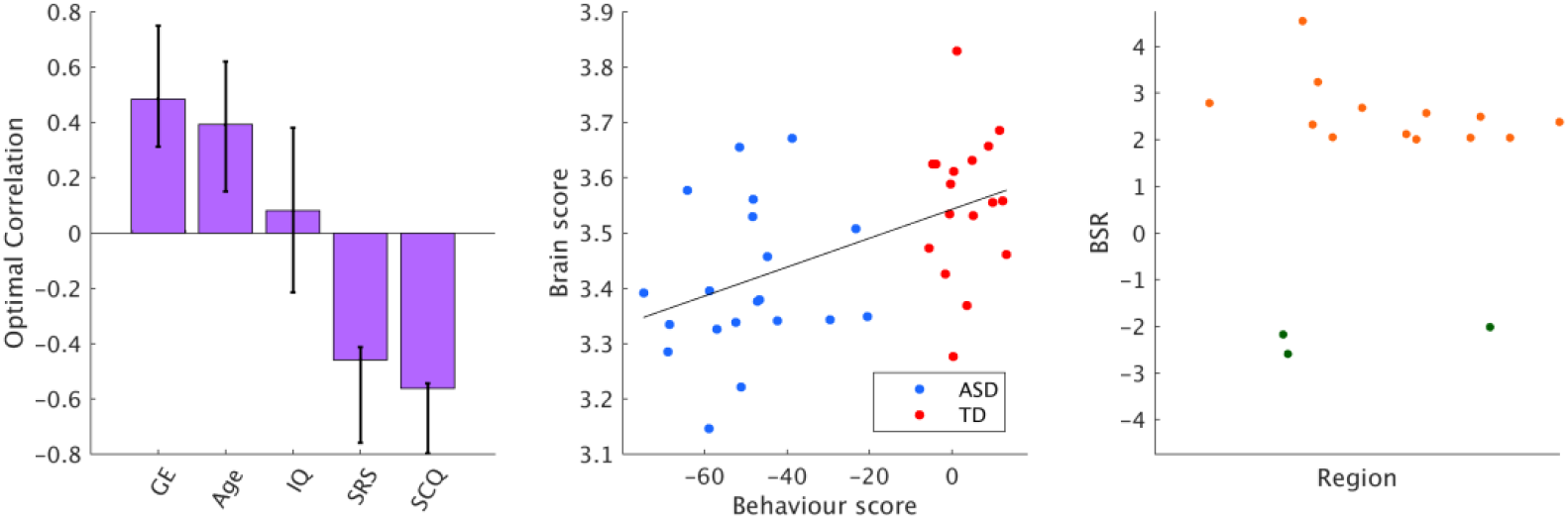
Contrast in relationships for correlations between entropy (averaged across m=2 conditions) and predictor variables, associated brain and behaviour scores for each group, and BSRs for each region, at a threshold of +2. Error bars show 95% confidence intervals determined through bootstrap resampling. Blue circles = ASD, red circles = TD.

**Figure 7:**
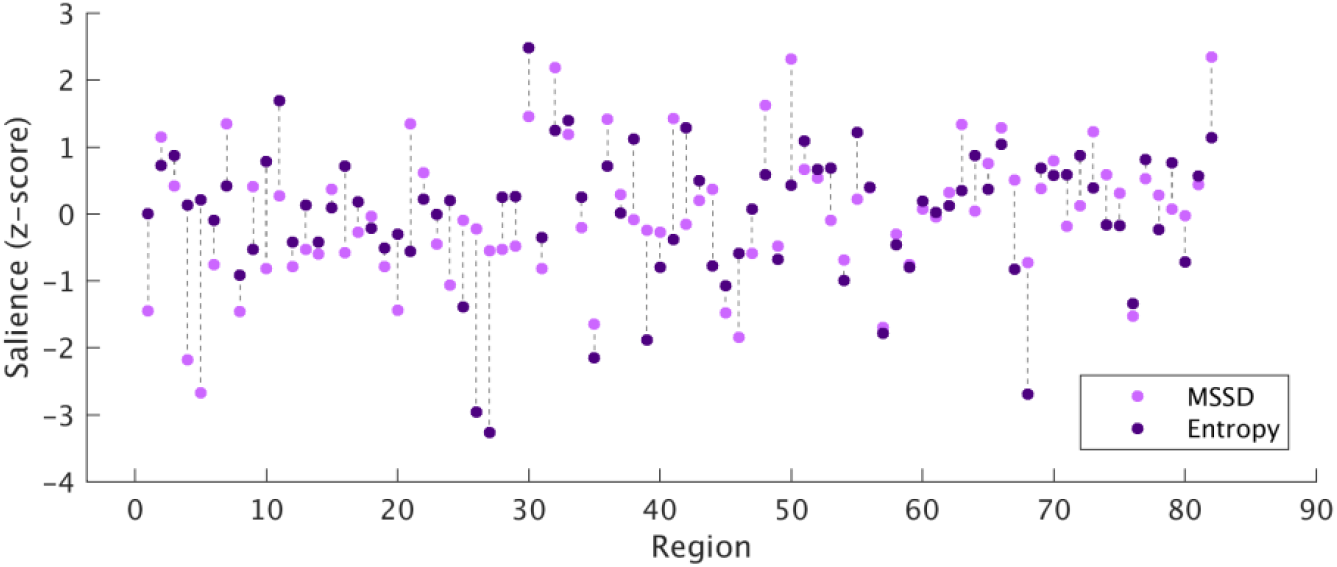
Brain saliences for MSSD and entropy. Light purple circles = MSSD, dark purple circles = entropy.

Another notable finding is that for both MSSD and entropy, behaviour scores were qualitatively more variable for the ASD group compared to the TD group, which illustrates the highly heterogeneous nature of ASD: for MSSD, SD_ASD_ = 12.33, and SD_TD_ = 5.09, *F*(19, 16) = 5.88, *p* < 0.001; for entropy, SD_ASD_ = 14.43, SD_TD_ = 6.10, *F*(19, 16) = 5.60, *p* = 0.001.

PLS was performed using SRS total scores as opposed to scores on the five SRS subscales, because scores on the subscales were highly correlated with each other as well as total SRS scores (*r* ≥ 0.80) and therefore biased the PLS results.

Behavioural PLS analysis was also performed in the ASD group using ADOS2 scores in addition to the other predictor variables. The correlation matrix for this set of variables for the ASD group is shown in Figure 8. As ADOS2 scores were only available for the ASD group, the TD group was not included in these analyses. When ADOS2 total scores were included, the PLS analysis was significant for MSSD (*p* = 0.02), but not for entropy (*p* = 0.17). However, in both cases, the behaviour saliences were not reliable for ADOS2 total scores, as the 95% CIs crossed the x-axis. A similar pattern was observed when the ADOS2 social affect (SA) and RRB scores were used instead of the total scores: PLS was significant for MSSD (*p* = 0.02) and not significant for entropy (*p* = 0.18), but the behaviour saliences for ADOS2 SA and RRB scores were not reliable.

**Figure 8:**
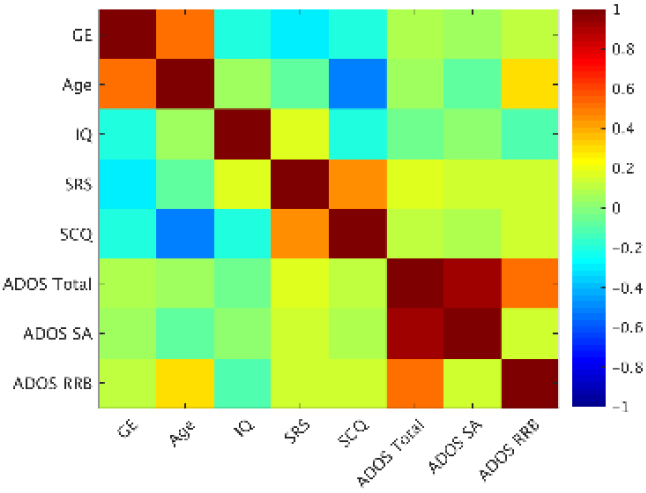
Correlation matrix for the set of behaviour variables used in the behavioural PLS analysis for the ASD group.

## Discussion

### Overview

Variability and complexity of resting-state BOLD signals were examined in participants with and without ASD. For both MSSD and entropy, a continuum of brain-behaviour relationships was observed across diagnostic groups despite a lack of significant differences between groups. A distributed set of brain regions exhibited positive correlations between MSSD and GE and age in both groups, and negative correlations with SRS and SCQ scores. A similar pattern was observed for entropy.

### Variability and complexity in ASD

Categorical and dimensional approaches were used to characterize MSSD and entropy in participants with ASD compared to TD participants. The categorical approach involved analyzing differences between diagnostic groups (ASD and controls), whereas the dimensional approach involved the use of continuous measures: age, IQ, SRS and SCQ scores, and GE. No group differences were observed when a categorical approach was used; however, using the dimensional approach, a set of brain regions exhibited relationships with the predictor variables.

The negative relationship between entropy and SRS and SCQ scores supports the notion that greater severity of ASD behaviours is associated with decreased entropy, and therefore decreased information processing capacity, in the brain. Few studies have examined entropy in ASD; however, our results are in line with a previous EEG study that found reduced MSE in ASD during both social and non-social tasks (Catarino et al., 2011). Reduced entropy in participants with more severe ASD behaviours supports the “loss of brain complexity hypothesis” proposed by Yang & Tsai (2013), which states that the complexity of neural signals reflects the capability of the brain to adapt to changing environments, and that in pathological states, this “adaptive capacity” of the brain is reduced.

Similarly, a continuum of brain-behaviour relationships was observed for MSSD, whereby SRS and SCQ scores were negatively correlated with MSSD. A dimensional approach has been used to study MSSD in ADHD. Nomi et al. (2018) did not find significant differences in MSSD between children with and without ADHD, but found positive correlations between MSSD and ADHD behavioural severity across diagnostic categories. While Nomi et al. (2018) found positive correlations between MSSD and symptom severity, we found negative correlations between these measures. This finding may seem counterintuitive to theories suggesting that ASD is characterized by an increased ratio of excitatory-inhibitory coupling in the brain, which may lead to poor functional differentiation and noisier and unstable neural signaling, resulting in inefficient information processing (Rubenstein & Merzenich, 2003). One possible explanation for this discrepancy is that detrimentally increased levels of noise may manifest at smaller scales in ASD, but not at macroscopic scales as measured by fMRI. This hypothesis is in line with the theory that physical connectivity at the level of microcolumns is increased in ASD, but computational connectivity is reduced (Belmonte et al., 2004). In other words, within neural assemblies, connections between synapses and fiber tracts are increased, but long-range connections between functional brain regions are reduced. The authors hypothesized that increased physical connectivity could lead to undifferentiated neural regions, which would preclude the development of effective communication between long-range functional regions. Thus, at macroscopic scales, ASD-like symptoms may be associated with reduced variability, which could be associated with a reduced capacity to explore different functional network configurations. Further, we included a larger set of predictor variables in our study compared to Nomi et al.’s study. In addition to behavioural severity, we included age and a measure of information processing in structural networks (GE). This analysis revealed important interactions between GE, age, and behavioural severity, whereby a distributed set of brain regions exhibited positive correlations with GE and age, but negative correlations with behavioural severity. Thus, factors such as the efficiency of structural networks and age may be modulating the relationship between MSSD, entropy and measures of behavioural severity.

### Variability and entropy increase with age

Distributed brain regions showed increases in MSSD and entropy from childhood through adolescence, suggesting that across development, information processing capacity increases in the brain (Figures 5 and 6). Previous work has suggested that increased information processing capacity over development results from a greater quantity of possible configurations of functional networks (McIntosh et al., 2010). Entropy of EEG signals has been shown to increase from childhood to adulthood, and is associated with higher accuracy and less variable reaction times during task performance (McIntosh et al., 2008; Lippé et al., 2009; Misic et al., 2010). Increases in entropy across development are thought to reflect increased integration and segregation in the brain (McIntosh et al., 2008; Garrett et al., 2013).

While developmental changes in variability and entropy have mostly been studied using EEG, Nomi et al. (2017) used fMRI to study changes in variability across the lifespan. In this study, central executive, default mode, visual, sensorimotor, and subcortical regions exhibited linear decreases in MSSD from ages 6 to 85, whereas nodes in the salience network and bilateral ventral temporal cortices exhibited increases in variability. The discrepancies between these findings and ours may relate, in part, to differences in age ranges and preprocessing strategies, which can affect estimates of BOLD signal variability (Garrett et al., 2010; 2013). Additional research on developmental changes in BOLD signal variability across different age ranges using various preprocessing strategies is required to resolve these discrepancies.

### Variability and entropy are associated with global efficiency in structural networks

Behavioural PLS also revealed positive correlations between MSSD of BOLD time series and GE in structural networks, and between entropy and GE (Figures 5 and 6). This finding supports the notion that structure shapes function in the brain (Honey et al., 2007, 2009, 2010; Deco et al., 2013). Further, there was a contrast in the relationship between GE and behavioural severity, such that brain regions that exhibited positive correlations with GE exhibited negative correlations with behavioural severity. Previous work has shown relationships between GE in structural networks and cognitive abilities. Berlot et al. (2016) showed that mild cognitive impairment (MCI) patients exhibited reduced GE of brain networks, which was associated with reduced cognitive control. Further, in Alzheimer’s disease, reduced GE has been associated with memory and executive function abilities (Reijmer et al., 2013). Collectively, these results suggest an important relationship information processing capacity in structural and functional networks and cognitive functioning.

### Limitations

There are several limitations of the current study. First, our sample size was small due to the limited number of high quality datasets with both DTI and resting-state fMRI data from the ABIDE II database. We chose to analyze data from a single site for this study. While previous work has demonstrated the feasibility of multisite DTI studies based on high concordance of fractional anisotropy and mean diffusivity measurements across five different scanning sites (Fox et al., 2012), the reliability of measurements of resting-state FC across scanning sites is less clear. FC measurements can be affected by differences in scanner manufacturers and acquisition protocols (Shinohara et al., 2017; Yu et al., 2018). Dansereau et al. (2017) analyzed multisite resting-state FC data across 8 scanning sites, and reported small to moderate between-site effects. Jovicich et al. (2016) studied difference in DMN FC in 5 healthy elderly participants who were scanned at 13 different sites with harmonized protocols, and found significant differences in temporal signal-to-noise ratio between sites. These differences were hypothesized to relate to differences in hardware and pulse sequences between sites. Nielsen et al. (2013) found that classification of ASD compared to controls based on resting-state FC was lower when multiple sites were included as opposed to a single site. Further, Easson et al. (2018) defined subtypes of ASD and controls based on FC using 5 sites from the ABIDE database, and found that it was necessary to regress out the effects of scan site prior to k-means classification, otherwise, the clusters differed significantly in scan site. This finding shows that prominent FC clusters defined in an unsupervised manner are driven by differences in FC across scanning sites. Thus, it will be crucial to further elucidate the effects of scanning site on resting-state FC to allow for multisite studies, and thus, larger sample sizes.

### Conclusions

This study reveals relationships between brain signal variability and complexity and GE, age, and behavioural severity across ASD and TD participants, and illustrates the importance of taking a dimensional approach to studying brain function in ASD. By analyzing brain variability and complexity in relation to a set of predictor variables, a continuum of relationships between brain variables and predictor variables was observed; however, when treating ASD and TD groups categorically, significant group differences were not observed. Further, increased GE in structural networks and higher age were associated with higher MSSD and entropy, revealing important information about structure-function relationships in the brain and the developmental trajectories of variability and complexity.

## Acknowledgements

This work is supported by an Ontario Graduate Scholarship and Finkler Graduate Student Fellowship to A.K. Easson, and a Natural Sciences and Engineering Research Council of Canada (NSERC) grant (RGPIN-2018-04457) to A.R. McIntosh.

## Abbreviations

ABIDE: Autism Brain Imaging Data Exchange
ADOS2: Autism Diagnostic Observation Schedule 2
ASD: autism spectrum disorder
BSR: bootstrap ratio
GE: global efficiency
MSSD: mean square successive difference
PLS: partial least squares
RRB: restricted and repetitive behaviour
SA: social affect
SC: structural connectivity
SCQ: Social Communication Questionnaire
SRS: Social Responsiveness Scale
TD: typically developing

## Notes

**Conflict of interest:** The authors declare no competing financial interests.

